# Targeting Metabolic Vulnerabilities: Valinomycin Augments the Potency of Bioenergetic Inhibitors in Combatting Drug-Resistant and Dormant *Mycobacterium tuberculosis*

**DOI:** 10.1101/2025.11.23.690070

**Authors:** Arnab Roy, Satish Kushwah, Shashikanta Sau, Puja Kumari Agnivesh, Kishan Kumar Parida, Madhe Divya, Nitin Pal Kalia

## Abstract

Tuberculosis (TB) treatment is hampered by monotherapy limitations and phenotypic drug tolerance, citing the need for effective drug combinations. The mycobacterial electron transport chain (ETC), crucial for oxidative phosphorylation and ATP production in dormant *Mycobacterium tuberculosis* (*Mtb*), is a key target. This study investigates combining established bioenergetic inhibitors-bedaquiline (BDQ), telacebec (Q203), and clofazimine (CFZ) with the potassium ionophore valinomycin, which disrupts the proton motive force (*pmf*). We demonstrated that valinomycin significantly potentiated the anti-TB activity of these inhibitors against both replicating and nutrient-starved non-replicating *Mtb*. Checkerboard assays revealed synergistic activity with BDQ and additive effects with Q203 and CFZ, correlated with a two to three-fold reduction in ATP IC_50_ values. Critically, valinomycin converted the bacteriostatic activity of the inhibitors at sub-MIC concentration into bactericidal killing in time-dependent killing assay, achieving sterilization. This lethal synergy for the combinations was also observed in a THP-1 macrophage intracellular model for TB. Respiration assays confirmed that the combinations collectively halted oxygen consumption. We conclude that concurrently targeting specific ETC on *Mtb*’s bioenergetics. This strategy, particularly the BDQ/valinomycin synergy, represents a promising cornerstone for developing novel sterilizing regimens to shorten TB therapy and overcoming drug tolerance.

## Introduction

Serious issues regarding global health are raised by the evolution and spread of antibiotic resistance in pathogenic mycobacteria. Global tuberculosis report 2025 stated that a total of 1.23 million people died from tuberculosis (TB) in 2024 making it the 2^nd^ leading cause of death from a single infectious agent (World Health Organization, 2025). 27% of the TB cases are from India suggesting it to be the country with highest burden of the disease. There is an ongoing increase in the prevalence of multi drug-resistant (MDR) and extensively drug-resistant (XDR) TB infections, even with advancements in healthcare administration and the implementation of fixed-dose regimens. An estimated 390,000 people were diagnosed with MDR-TB in 2024 and treatment is difficult due to the requirement of administering second line anti-TB drugs for two years (World Health Organization, 2025). The creation of novel medications with the potential to reduce the duration of MDR-TB treatment to six months or less is an urgent therapeutic necessity (Conradie et al., 2020). More than novel medications, we desperately need a logical pharmacological combination consisting of complementary drugs (Dartois and Barry, 2013). The worldwide TB drug pipeline is still quite narrow, despite growing interest from the scientific community: in the last 40 years, very few novel molecules have entered the clinical phases of the drug discovery process (Ginsberg and Spigelman, 2007). Bedaquiline (BDQ, Sirturo) recent approval marks a significant advancement in the search for anti-TB drugs (Andries et al., 2005). However less than three years after BDQ was first used clinically, resistance cases began to emerge, overshadowing the drug’s remarkable advancement (Bloemberg et al., 2015). The lack of effective companion medications is probably a contributing factor to the quick evolution of resistance. Currently, BDQ is used in conjunction with less effective second- and third line drugs, which places a significant burden on BDQ resistance. This supports the idea that to reduce the duration of time needed to treat MDR-TB, a sensible pharmacological approach with effective drug combination is needed.

Oxidative phosphorylation (OxPhos) came in limelight as a lucrative drug target by the identification of BDQ, an effective inhibitor of the F_1_F_0_ ATP synthase in the mycobacterial electron transport chain (ETC) (Andries et al., 2005). OxPhos metabolic pathway is widely distributed and involves the utilization of nutritional energy to produce an electrochemical gradient, commonly known as the proton motive force (*pmf*), which facilitates the production of ATP (Hards and Cook, 2018). Both replicating and non-replicating phase (also known as dormant state) of mycobacteria depend on the *pmf* to survive. Cell death and a rapid loss of viability can be achieved by disruption of the *pmf* (Rao et al., 2008; Cook et al., 2014). Therefore, targeting the various components involved in pmf production can help shorten the duration of therapy by eliminating subpopulations of the bacteria that are phenotypically drug-resistant (Bald et al., 2017).

In recent years, bioenergetics has drawn a lot of attention as a promising field for the discovery of anti-TB drugs after the approval of bedaquiline in 2014. Targeting the ETC and OxPhos machinery particularly can eliminate both replicating as well as the metabolically quiescent bacilli, which are inherently more drug-tolerant (Koul et al., 2011). Bioenergetic inhibitors like BDQ, telacebec (Q203), and clofazimine (CFZ) target energy metabolism of *Mtb*, depleting bacterial energy (Cholo et al., 2012; Pethe et al., 2013). As demonstrated in this study, these effects can be potentiated by ionophores like valinomycin, which directly dissipates the ion gradient offering a powerful combinatorial strategy to combat against the core metabolism of *Mtb* (Rao et al., 2008; Hards et al., 2018).

## Results

### Valinomycin enhanced the growth inhibition efficacy of bioenergetic inhibitors

To quantify the combined inhibitory effect of valinomycin with bioenergetic inhibitors, we performed the checkerboard broth microdilution assays against *Mtb* mc^2^6230 and determined fractional inhibitory concentration (FIC) index. The growth inhibition assays revealed that sub-inhibitory concentrations of valinomycin markedly increased the potency of BDQ, Q203, and CFZ. The minimum inhibitory concentration leading to 50% growth inhibition (MIC_50_) for each drug was significantly reduced in a dose dependent manner with the presence of valinomycin, as evidenced by the shift in the RFU values in (Figures 1A, 1B, 1C). The interaction between BDQ and valinomycin was found to be synergistic, with an FIC index of 0.36 (Figure 1D). The MIC_50_ of BDQ was reduced from 75.01 nM to 11.86 nM (6.3-fold reduction), while the effective MIC_50_ of valinomycin decreased from 21.19 nM to 4.55 nM. The combinations of valinomycin with CFZ and Q203 were classified as additive, with FIC indices of 0.98 and 1.08 respectively. Notably, the combination with CFZ resulted in a dramatic 247-fold reduction in its effective MIC_50_, from 375.6 nM to 1.52 nM.

**Figure 1.**
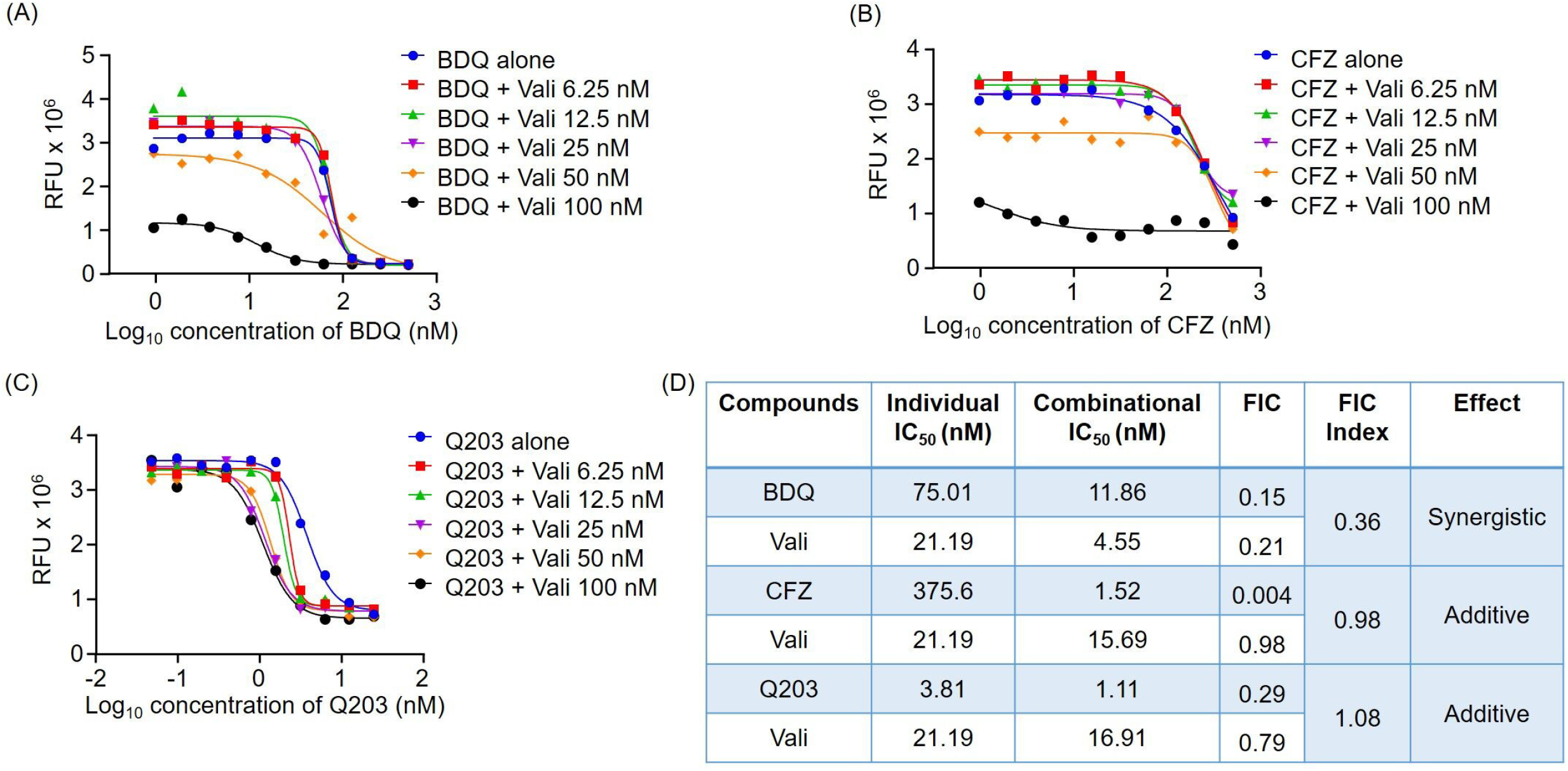
Valinomycin mediated enhanced efficacy of BDQ, CFZ, and Q203. (A) BDQ susceptibility in *Mtb* mc^2^6230 in combination with indicated concentrations of valinomycin. (B) Inhibitory effect of CFZ in *Mtb* mc^2^6230 in combination with indicated concentrations of valinomycin. (C) Valinomycin increases the potency of Q203 growth inhibition in *Mtb* mc^2^6230. (D) Table indicating the individual MIC_50_ values along with the combination MIC_50_ values of the indicated drug combinations. Calculated FIC and FICI values along with indicative effect is also represented, where BDQ and valinomycin combination shows synergistic effect and the other two bioenergetic inhibitors displays additive effect. Bacterial growth was measured by recording the relative fluorescence unit (RFU) at 530/590 nm excitation/emission after 10d of incubation and addition of resazurin.

These results demonstrate that valinomycin potently enhances the in vitro growth inhibitory activity of a range of bioenergetic inhibitors. The strong synergy with BDQ suggests that the simultaneous disruption of the membrane potential by valinomycin and inhibition of ATP synthase by BDQ creates a particularly lethal combination for the bacilli.

### Combination of valinomycin and bioenergetic inhibitors break the ATP homeostasis in both replicating and non-replicating *Mtb*

Having established that valinomycin synergistically enhances the growth inhibitory effects of bioenergetic inhibitors, we next sought to determine the impact of the combinations on bacterial viability. We employed an ATP depletion assay to measure the disruption of core bioenergetics in both replicating and non-replicating (nutrient starved) *Mtb* mc^2^6230. Against replicating mycobacteria, the combination of valinomycin with BDQ, CFZ, and Q203 consistently enhanced ATP depletion a concentration dependent manner (Figure 2A. 2B, 2C). Quantifying the data confirmed an additive interaction for all three combinations (Figure 2G). The BDQ/valinomycin combination was the most potent, reducing the ATP IC_50_ of BDQ from 112.5 nM to 32.91 nM (3.4-fold reduction) and yielding a FIC index of 0.81. Similarly additive effects were observed for CFZ and Q203 with FIC index 0.79 and 1.00 respectively. This demonstrates that dissipating the membrane potential while inhibiting respiration exacerbates the bioenergetic crisis in replicating mycobacteria. We then evaluated the same combinations against the phenotypically drug-tolerant, non-replicating population. Strikingly, the combinations remained effective, again demonstrating additive effects in depleting the basal ATP levels essential for survival of dormant mycobacteria (Figure 2D, 2E, 2F). The FIC indices for all the pairs were additive ranging in between 1.37 to 1.44 (Figure 2H). This indicates that even against non-replicating *Mtb*, where ATP synthase activity is downregulated, the dual targeting of the ETC and the membrane potential via valinomycin creates a bioenergetic disruption to compromise viability. Collectively, these findings confirm that the functional synergy observed in growth inhibition assays is linked with the disruption of the mycobacterial energy homeostasis. This consistency of this effect across both metabolic states underscores the vulnerability of mycobacteria to combined ionophore and bioenergetic inhibitor treatment and highlights a promising strategy to target hard to eradicate, non-replicating populations.

**Figure 2.**
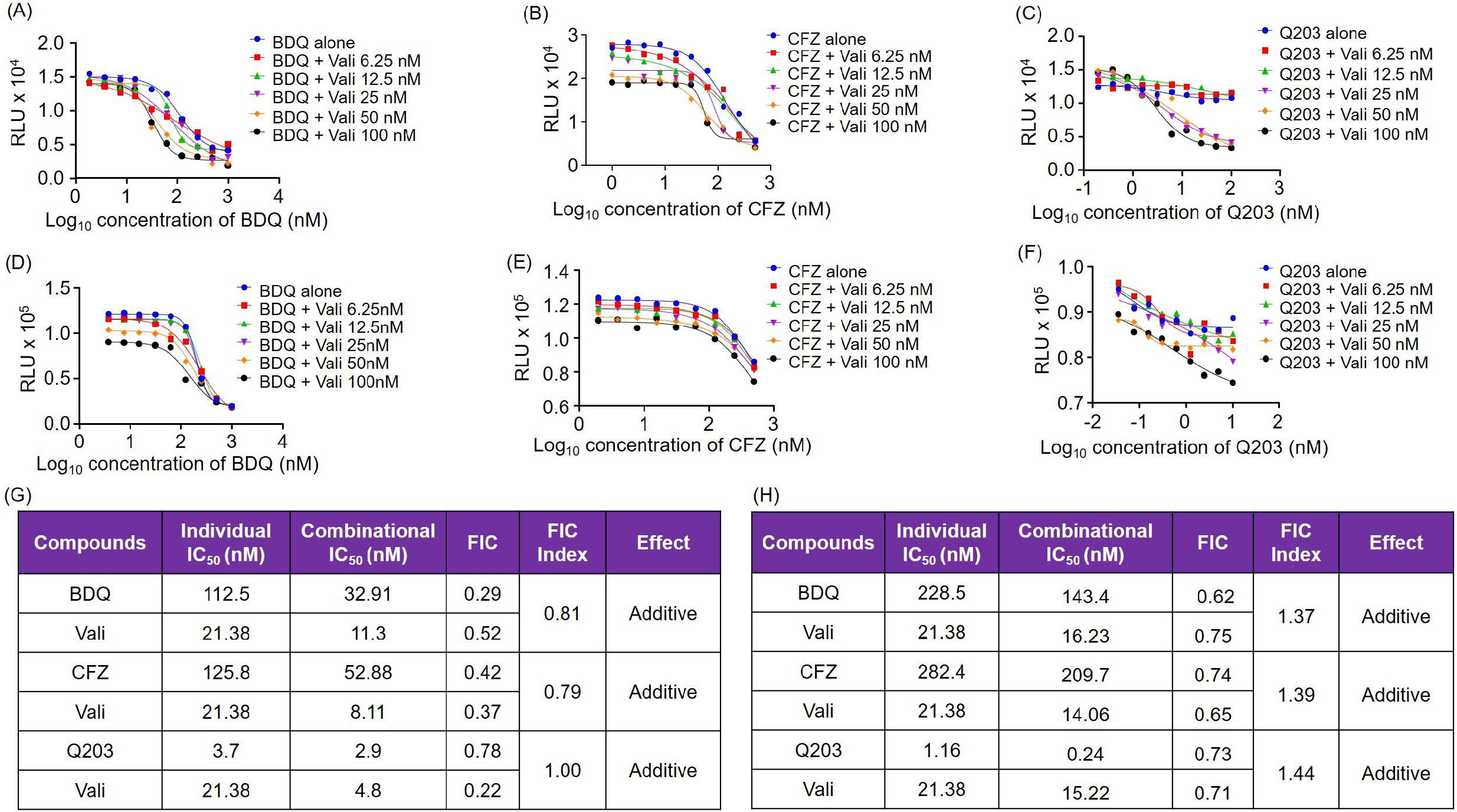
Valinomycin synergized with bioenergetic inhibitors and resulted in abrupt ATP depletion in mycobacteria under replication and non-replicating conditions. (A) and (C) Intracellular checkerboard ATP depletion assay for BDQ and valinomycin combination in replicating and non-replicating (nutrient starved) *Mtb* mc^2^6230 respectively. (B) and (D) Intracellular checkerboard ATP depletion assay for CFZ and valinomycin combination in replicating and non-replicating (nutrient starved) *Mtb* mc^2^6230 respectively. (C) and (E) Intracellular checkerboard ATP depletion assay for Q203 and valinomycin combination in replicating and non-replicating (nutrient starved) *Mtb* mc^2^6230 respectively. (F) Table indicating the individual ATP IC_50_ values along with the combination ATP IC_50_ values of the indicated drug combinations and the FIC index in *Mtb* mc^2^6230. (G) Table indicating the individual ATP IC_50_ values along with the combination ATP IC_50_ values of the indicated drug combinations and the FIC index in non-replicating (nutrient starved) *Mtb* mc^2^6230.

### Killing efficacy of bioenergetic inhibitors enhanced with valinomycin in heterogenous population of *Mtb*

The functional synergy observed in growth inhibitory and ATP depletion assays prompted an investigation into the bactericidal consequences of combining valinomycin with bioenergetic inhibitors. Against replicating *Mtb* mc^2^6230, monotherapy with BDQ, CFZ, and Q203 (each at 100 nM) or valinomycin (100 nM) proved bacteriostatic over a 10-day exposure, failing to reduce the bacterial load by 90% (MBC_90_). In sharp contrast, the combination valinomycin with any of the three inhibitors resulted in a profound bactericidal outcome, with the BDQ/valinomycin pair reducing viable counts to levels approaching the limit of detection (Figure 3A). This bactericidal effect proved critical against phenotypically resistant, nutrient starved (non-replicating) *Mtb* mc^2^6230. In this model, which mirrors the drug tolerant state responsible for protracted therapy, individual compounds were largely ineffective. However, the concurrent administration of valinomycin with each bioenergetic inhibitor restored potent killing activity, surpassing the MBC_90_ and demonstrating the necessity of targeting the *pmf* to eradicate dormant populations (Figure 3B). Time-dependent killing assay provided further insights into the killing efficacy of the combinations. The BDQ/valinomycin combination induced a rapid decline in the bacterial viability, resulting in complete sterilization by day 9. A similar, though temporarily delayed, sterilizing effect was observed with Q203/valinomycin, which achieved complete bacterial clearance by day 12 (Figure 3C). These data illustrate that the partnership not only enhances the endpoint efficacy but also dramatically accelerates the rate of kill. The combinations were also tested against the MDR strain *Mtb* mc^2^8259 and it proved to be equally potent (Figure 3D). This suggests that the combinations can be effective against the drug-resistant strains. The therapeutic potential of this strategy was validated intracellularly. In *M. smegmatis* infected THP-1 macrophages, the combinations of BDQ/valinomycin and Q203/valinomycin exhibited significantly enhanced bactericidal activity compared to their components alone, each achieving a >90% reduction in bacterial burden post-infection (Figure 3E). In summary, the addition of the ionophore valinomycin fundamentally alters the profile of bioenergetic inhibitors, converting them into a potent sterilizing regimen capable of targeting heterogenous subpopulations of mycobacteria.

**Figure 3.**
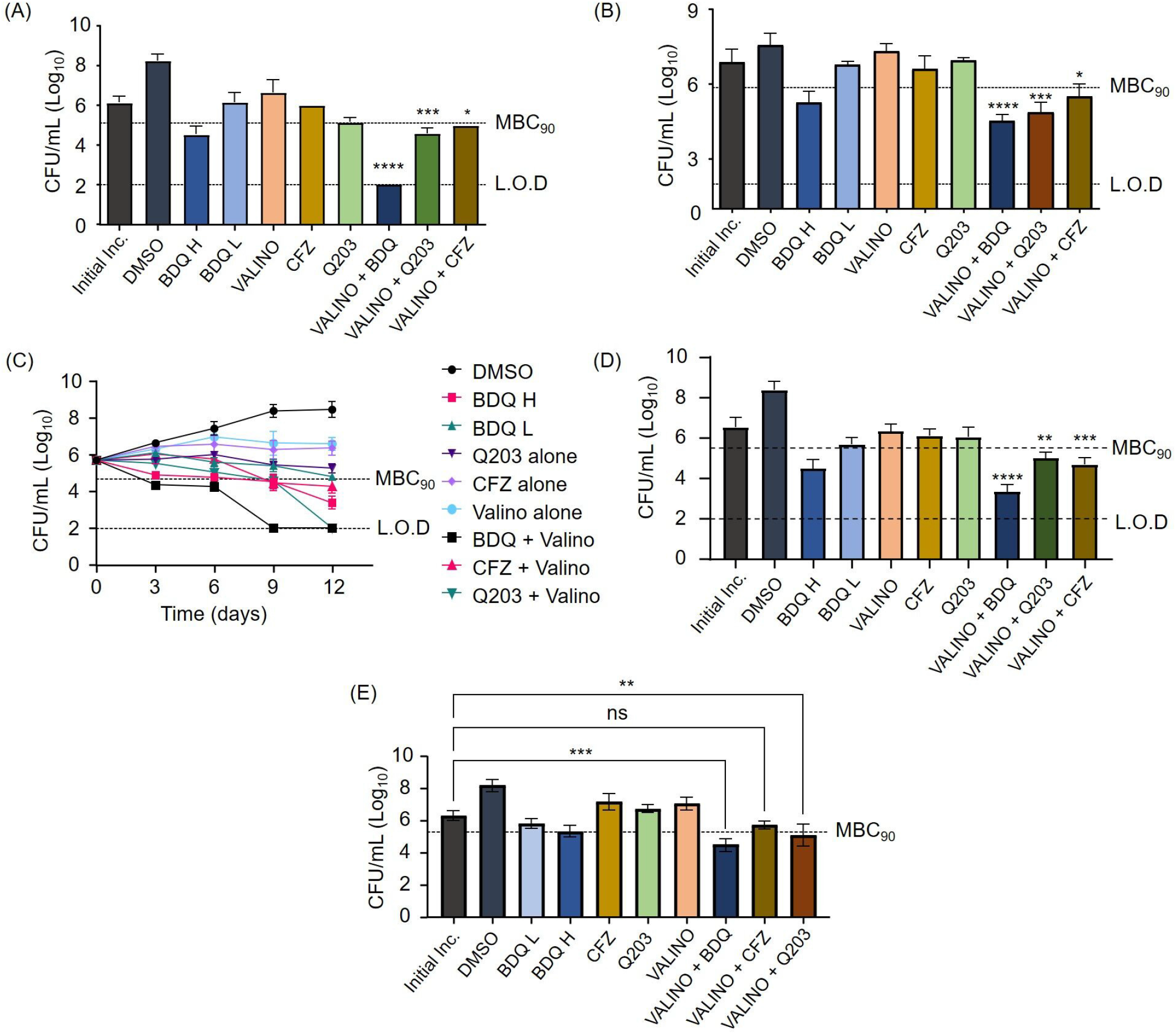
Valinomycin enhanced the killing efficacy of bioenergetic inhibitors against heterogenous population of *Mtb*. (A) Bactericidal activity of the bioenergetic inhibitors and valinomycin against replicating *Mtb* mc^2^6230. Viability of bacteria was determined by taking the CFU on 7H10 agar plates after 10d of incubation. (B) Bactericidal activity of the bioenergetic inhibitors and valinomycin against non-replicating (nutrient starved) *Mtb* mc^2^6230. Viability of bacteria was determined by taking the CFU on 7H10 agar plates after 15d of incubation. (C) Time-dependent killing efficacy of the drug combinations. CFU were taken in interval of 3 days up to 12 days of incubation period. (D) Bactericidal activity of the bioenergetic inhibitors and valinomycin against multi-drug resistant *Mtb* mc^2^8259. Viability of bacteria was determined by taking the CFU on 7H10 agar plates after 10d of incubation. (E) THP-1 cells were infected with *M. smegmatis* and treated with the drug combinations for 48 hours. Viability of intracellular bacteria was determined after 2d of treat post-infection. Concentrations of the inhibitors used: BDQ L- 100 nM, BDQ H-1 µM, CFZ-100 nM, Q203-100 nM, and valinomycin-100 nM. In Inc: starting inoculum, DMSO: vehicle control. Dotted line with MBC_90_ represents 90% bacterial killing, LOD represents limit of detection for the experiment.

### Valinomycin induces a collective arrest of mycobacterial respiration and altered membrane potential, reason for greater efficacy of the bioenergetic inhibitors

Given the central role of oxygen as the terminal electron acceptor in the mycobacterial ETC, we directly assessed the impact of our drug combinations on bacterial respiration using methylene blue as the oxygen probe. In this system, the persistence of blue colour indicates the presence of oxygen, while decolourization signifies oxygen depletion by respiring bacilli. As anticipated the, subinhibitory concentrations of the bioenergetic inhibitors when administered as monotherapies, were insufficient to halt mycobacterial respiration. In these vials, methylene blue was fully decolourized within 96-hour period (Figure 4A), confirming that the bacilli continued to consume oxygen. In contrast, the combination of each bioenergetic inhibitor with valinomycin resulted in collective suppression of oxygen consumption. The vials containing these combinations retained a distinct blue colour throughout the 96-hour observation window (Figure 4A). This qualitative result provides compelling evidence that the drug pairs act synergistically to induce a state of complete respiration arrest. The failure to deplete the environmental oxygen underscores the catastrophic breakdown in the electron flow, from the dual inhibition of enzyme complexes and the dissipation of *pmf* by the ionophore. This collective respiratory collapse provides a mechanistic basis for observed bactericidal synergy and ATP depletion, confirming that the combinations fundamentally disrupt a core physiological process essential for mycobacterial survival.

**Figure 4.**
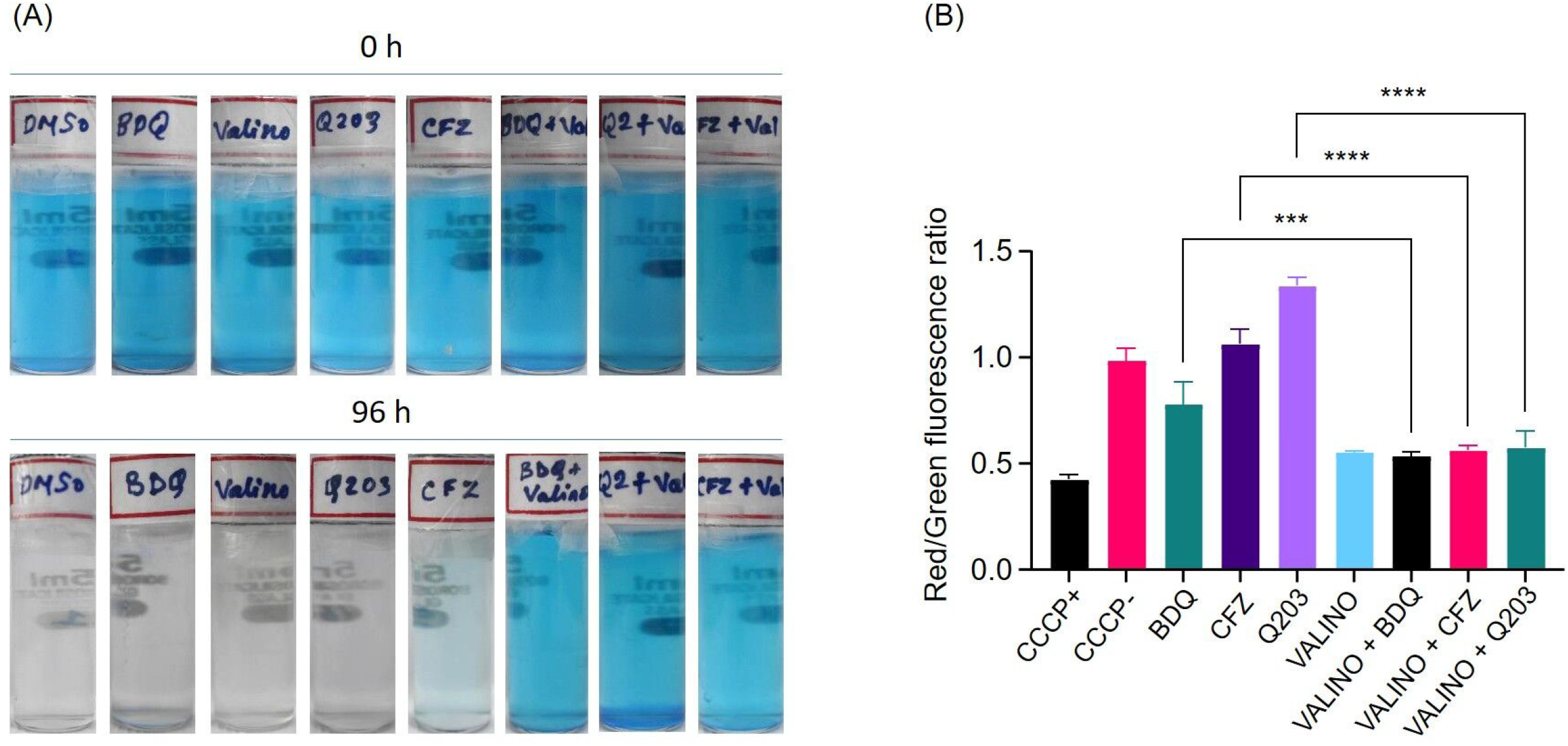
Alteration in membrane potential and complete respiration inhibition by BDQ, CFZ, and Q203 in presence of valinomycin, reason for enhanced efficacy against *Mtb*. (A) Oxygen consumption assay in *Mtb* mc^2^6230 using methylene blue as an oxygen sensor probe at 0.001%. (B) Effect of the drug combinations with valinomycin on the membrane potential of *Mtb* mc^2^6230. Results are represented as Red/Green fluorescence ratio after staining with DiOC_2_.

The *pmf*, a cornerstone of mycobacterial bioenergetics, comprises a proton gradient and membrane potential (ΔΨ). Given that valinomycin is a potassium ionophore known to dissipate ΔΨ, we sought to quantify the impact of our drug combinations on the membrane potential of *Mtb*. To test this, we directly assessed the integrity of the ΔΨ using a fluorescent dye DiOC_2_. As anticipated, valinomycin alone, a known potassium ionophore, effectively collapsed the ΔΨ, resulting in a significantly diminished fluorescence ratio. In contrast, the bioenergetic inhibitors BDQ, Q203 and CFZ, when analyzed as monotherapy, induced only a marginal depolarization, indicating that their primary action on specific enzyme complexes does not directly trigger a full collapse of the membrane potential. Critically, the combinations of valinomycin with each bioenergetic inhibitor produced the most profound effect. The simultaneous use of the ionophore and a respiratory inhibitor led to near-complete dissipation of the ΔΨ, with the fluorescence ratio dropping to levels significantly below those of any single drug treatment (Figure 4B). This demonstrates that the collective action of disrupting electron flow and directly shunting ionic gradients synergizes to cripple the mycobacterial membrane potential. This provides a direct mechanistic explanation for the observed bactericidal and ATP depletion, positioning the collapse of ΔΨ as the central event in the synergistic killing of *Mtb*.

## Discussion

The limitations of monotherapy for tuberculosis, particularly the propensity for resistance development, underscore the critical need for exploring effective drug combinations (Nuermberger and Grosset, 2004). This is especially pertinent for targeting phenotypically drug-tolerant, non-replicating populations of *Mtb*. The mycobacterial ETC is a validated target in this context, as it is indispensable for OxPhos and ATP generation, processes upon which *Mtb* relies during dormancy (Gengenbacher and Kaufmann, 2012; Cook et al., 2014). The last decade has seen the development of several bioenergetic inhibitors, such as the ATP synthase inhibitor BDQ and the cytochrome *bc*_*1*_*:aa*_*3*_ inhibitor Q203 (Andries et al., 2005; Pethe et al., 2013). Furthermore, the increased ETC activity induced by some agents can elevate reactive oxygen species production, a vulnerability exploited by drugs like CFZ (Yano et al., 2011). Here we demonstrated that the potassium ionophore valinomycin significantly potentiates the efficacy of these bioenergetic inhibitors by disrupting the *pmf*, revealing a promising combinatorial strategy.

Our initial findings derived from checkerboard broth microdilution assays against replicating *Mtb*, confirmed that valinomycin enhances the inhibitory effects of BDQ, Q203 and CFZ. The combination of BDQ and valinomycin was unequivocally synergistic while combinations with Q203 and CFZ were additive. This potentiation was corroborated by the ATP depletion assay. We observed a two to three-fold reduction in the ATP IC_50_ for all three combinations, indicating a heightened disruption of bacterial bioenergetics. Given that ATP synthase utilizes the *pmf*-comprising a ΔΨ component for ATP production, the additive effect likely stems from the dual inhibition of both specific ETC components by the inhibitors and the Δψ by valinomycin (Hards and Cook, 2018; Hards et al., 2018). This mechanism is particularly critical for non-replicating mycobacteria, which maintain a reduced but essential level of ATP and require an energized membrane for survival (Rao et al., 2008). Indeed, against nutrient-starved, non-replicating *Mtb*, the combinations yielded a two-fold reduction in ATP IC_50_, demonstrating that bioenergetic inhibitors coupled with an ionophore effectively exploit the metabolic vulnerability of quiescent bacilli by crippling ATP homeostasis.

To further characterize the therapeutic potential of these combinations, we performed MBC and time-kill kinetic assays. As previously reported, BDQ, Q203 and CFZ individually exhibited a bacteriostatic phenotype against replicating *Mtb* under our conditions (Koul et al., 2014; Kalia et al., 2017). Notably the presence of valinomycin converted these static responses into bactericidal activity, achieving a 90% reduction in bacterial load within 10 days. The combination of BDQ and valinomycin led to complete sterilization by day 9, while Q203 with valinomycin achieved the same effect by day 12. This suggests that dissipating the Δψ with an ionophore is a critical determinant for promoting lethal synergy when core respiration is inhibited. This principle was further validated in a nutrient-starved non-replicating model, where the bioenergetic inhibitors alone at low concentrations were ineffective, but all combinations with valinomycin were bactericidal. This finding confirms the importance of simultaneously targeting respiratory components and the *pmf* to create a sterilizing regimen against phenotypically drug-tolerant populations. The translational relevance of these findings was assessed using THP-1 macrophage infection model. The combinations of BDQ/valinomycin and Q203/valinomycin demonstrated significantly enhanced bactericidal activity compared to individual drugs, underscoring their efficacy within a host-mimicking environment. The CFZ/valinomycin combination, however, showed the least bactericidal activity in this model, warranting further investigations into its specific intracellular pharmacokinetics.

Finally, to directly visualize the impact on bacterial respiration we employed an oxygen consumption assay using methylene blue as an indicator (Lee et al., 2021). While low concentrations of the bioenergetic inhibitors alone failed to prevent oxygen depletion (evidenced by discolouration of methylene blue), all three combinations with valinomycin abruptly halted oxygen consumption, with the dye retaining its blue colour after 96 hours. This qualitative result provides direct visual correlation with our other findings, confirming that the combinations induce a profound and collective arrest of mycobacterial respiration. Our findings provide a direct basis for the observed synergistic and bactericidal effects of combining valinomycin with bioenergetic inhibitors. While BDQ, Q203, and CFZ as monotherapies caused only marginal depolarization, their combination with valinomycin resulted in a near-complete collapse of the ΔΨ. This demonstrates that the primary action of these inhibitors on specific ETC complexes is insufficient to fully dissipate the membrane potential. The profound depolarization achieved by the combinations arises from a two-way attack: valinomycin directly shunts the ionic gradient, while the bioenergetic inhibitors prevent the ETC from compensating for this dissipation. We therefore, can say that the collapse of ΔΨ is the central mechanistic event driving the potent bactericidal activity and ATP depletion, underscoring the critical importance of simultaneously targeting respiratory function and the membrane’s energetic integrity.

In conclusion, our study, establishes that combining bioenergetic inhibitors with the ionophore valinomycin creates a powerful, multi-pronged attack on *Mtb*’s central energy metabolism. By concurrently inhibiting specific ETC complexes and dissipating the essential membrane potential, these combinations effectively kill heterogenous population of mycobacteria. This approach, particularly the synergistic pairing of bedaquiline and valinomycin, represents a cornerstone strategy for developing novel sterilizing regimen aimed at shortening TB therapy and overcoming phenotypic drug tolerance.

## Materials and Methods

### Bacterial strains and Culture conditions

*Mtb* auxotrophic strain mc^2^6230 was cultivated in Middlebrook 7H9 broth. The medium was supplemented with 10% albumin-dextrose-catalase (ADC), 0.05% Tween 80, 0.2% glycerol, L-methionine (50 mg/L), L-arginine (200 mg/L), L-leucine (50 mg/L), and D-pantothenic acid (24 mg/L). All cultures were incubated at 37°C with 5% CO_2_. For drug susceptibility testing, glycerol was omitted from the medium. The human monocytic THP-1 cell line was maintained in RPMI 1640 medium, supplemented with 10% fetal bovine serum (FBS), 2 mM L-glutamine, and 10 mM sodium pyruvate (Vilchèze et al., 2018).

### Determination of Minimum Inhibitory and Bactericidal concentrations

MIC_50_ was determined from the checkerboard broth microdilution method in 96-well flat bottom plates. The MIC_50_ was defined as the lowest drug concentration that resulted in 50% reduction in bacterial growth compared to the untreated control (Kumar et al., 2025).

To determine the minimum bactericidal concentration for 90% killing (MBC_90_), bacteria at an initial OD_600_ of 0.005 were exposed to drugs for 10 days (replicating) or 15 days (non-replicating) at 37°C. Post incubation, bacterial viability was assessed by plating serial dilutions on Middlebrook 7H10 agar plates and enumerating colony forming units (CFU). The MBC_90_ was defined as the lowest drug concentration that reduced the initial bacterial inoculum by 90% (Chilakala et al., 2024).

### Intracellular ATP quantification

Bacterial ATP levels were measured using a luminescence-based assay. Drug combinations were prepared in a checkerboard format in 96-well white plates and inoculated with mycobacterial culture. After a 15-hour incubation at 37°C, the BacTiter-Glo™ microbial cell viability reagent (Promega) was added to each well. Following a 12-minute incubation in the dark, luminescence was recorded using a BioTek Cytation 5 multi-mode reader (Lee et al., 2021).

### Induction of nutrient starved model

Exponential-phase *Mtb* mc^2^6230 cultures were harvested by centrifugation washed with DPBS (containing Ca^2+^, Mg^2+^, and 0.025% Tween 80), and resuspended in the same media to an OD_600_ of 0.15. These cultures were then incubated for 2 weeks at 37°C prior to susceptibility testing (Kalia et al., 2017).

### Methylene blue assay

Bacterial respiration was qualitatively monitored using methylene blue as an oxygen probe. *Mtb* mc^2^6230 cultures (OD_600_=0.3) were pre-incubated with drugs for 4 hours in a sealed 5.5 mL glass tubes. Methylene blue was then added to a final concentration of 0.001%. The tubes were tightly sealed with parafilm and incubated for 96 hours in a lock and lock box containing anaerobic sachets (Saha et al., 2024).

### Intracellular THP-1 infection model

The intracellular efficacy of the drug combinations was evaluated using THP-1 macrophage infection model. Briefly, THP-1 cells were seeded in a 24-well plate at a density of 3×10^5^ cells per well and differentiated using 200 nM phorbol myristate acetate (PMA) for 24 hours. The differentiated monolayers were then infected with *M. smegmatis* at a multiplicity of infection (MOI) of 10 for 4 hours. Following infection, extracellular bacteria were removed by washing with pre-warmed PBS. The infected macrophages were subsequently incubated in RPMI 1640 medium containing the respective drug combinations. After 48 hours of treatment, the macrophages were lysed, and the intracellular bacterial viability was quantified by plating serial dilutions of the lysates on Middlebrook 7H10 agar for CFU determination (Riaz et al., 2020).

## Author Contribution

N.P.K designed research; A.R., S.K, S.S., P.K.A., K.K.P., and M.D., performed research; A.R, and N.P.K analyzed data; A.R. and N.P.K. wrote the paper; and all authors contributed to writing the paper.

## Acknowledgement

We thank Prof. William R. Jacob, <u>Albert Einstein College of Medicine</u> for the gift of the derivative strains of *M. tuberculosis* H37Rv and we are also thank Prof. Kevin Pethe for Providing Q203 and BDQ. This research is supported by the Indian Council of Medical Research (ICMR) New Delhi, India under its Investigator-Initiated Research Proposals - Small Grant (Project Award EM/Dev/SG/171/0227/2023), Department of Biotechnology, India (<u>D.O.NO.BT/HRD/35/02/2006</u>; NO. BT/RLF/Re-entry 66/2017), and National Institute of Pharmaceutical Education and Research Hyderabad under the Department of Pharmaceuticals, Ministry of Chemicals and Fertilizers, Govt of India.

